# Interconnection between molecular regulators in LMNA related muscular dystrophy

**DOI:** 10.1101/2022.03.09.483617

**Authors:** Subarna Dutta, Madavan Vasudevan

**Affiliations:** Department of Biochemistry,University of Calcutta, 35, Ballygunge circular road, Kolkata-700019.West Bengal. India; Theomics International Private Limited.28, Income Tax Layout, Sadananda Nagar, NGEF Layout, Bengaluru – 560038.India

## Abstract

Nuclear lamina is composed of different A-type and B-type lamin proteins and acts as major regulator of DNA replication, transcription, heterochromatin-euchromatin machinery. The majority of mutations in A type lamin are associated with some forms of muscular dystrophies,that majorly distinguishable by a common clinical feature : progressive skeletal muscle wasting. This speculates impaired skeletal muscle differentiation during development and after injury with the influence of myriad of signalling pathways. The molecular mechanism behind phenotypes of lamin A associated muscular dystrophies are still elusive . Here, we used genome wide expression analysis platform during the differentiation of murine C2C12 skeletal muscle cells to investigate the factors have been attributed to the disease. We compared the expression levels of the components of pathways that indicates important role of skeletal muscle as well as identified gene regulatory networks at two different time points of muscle differentiation in wild type and mutant cells. We also report significant perturbations in the expression and activation of Wnt signalling pathway in mutant cells among 40 dysregulated signalling pathways , presenting pronounced regulation of normal downstream myogenic signalling . Finally, with this largest data sets we evaluate in depth characterization of molecular effectors for myogenic differentiation, which could allow greater insight into development of therapeutic strategies for the remission of patients with LMNA linked muscle - related pathologies.

## Introduction

Skeletal muscle fibers are made up of bundles of multinucleated myotubes with actin-myosin contractile filaments that make up 40%-45 % of total body mass (Kwee, Mooney.2017). Myogenesis potentiality is required for skeletal muscle repair or regeneration (Tedesco et al. 2010). During muscle injury, a separate pool of muscle progenitor cells, known as satellite cells, which sit underneath the basal lamina, activate their myogenic potential and produce sufficient numbers of myotubes to quickly repair muscle damage. The process of myogenesis is controlled by distinct genetic networks. Members of the MyoD family of myogenic regulatory factors (MRFs) are expressed in chronological fashion and operate as key determinants during the muscular differentiation process. MyoD (Pinney et al. 1988), myogenin (Wright et al. 1989), Myf5 (Braun et al. 1989), and MRF4 (Braun et al. 1990) are the proteins in this event. Any of these factors can be induced to promote myoblast fusion by activating the myogenic potential in non-muscle cells (Davis et al.1987, Lattanzi et al. 1998). A combination of signaling molecules also causes MRFs to express directly for differentiation (Bryson-Richardson et al. 2008). Wnts, Sonic, Hedgehog and Noggin signaling pathways all play a role in MRF induction (Tajbakhsh et al. 1996, Borello et al. 1999). FGF and BMP4 signaling pathways, on the other hand, have been found to prevent satellite cell activation and differentiation (Itoh et al. 1998, Milasincic et al. 1996, Tajbakhsh et al. 1996). During myogenesis, Iwasa et al. 2003 observed activation of the PI3K and p38 MAPK signaling pathways, as well as inhibition of the Raf-MEK-ERK kinase pathways. The activation of the PI3K-AKT kinase signaling cascade by insulin growth factor (IGF) was discovered to trigger myogenesis (Orzechowski et al. 2005, Favreau et al.2008). Some epigenetic regulators, such as histone acetyltransferases, histone deacetylase (HDAC), and SWI/SNF chromatin remodelers, are also required for MRF activation (Naya et al. 1999, Puri et al. 2001). Gillespie et al. 2009 discovered that the p38 gamma signaling pathway enhances MyoD’s interaction with histone methyltransferase, which provides H3K9 methylation. This finding shows that in the differentiation process, there may be a relationship between signaling pathways and epigenetic mechanisms.

The LMNA gene has more than 500 mutations and is linked to 16 disorders, including Emery Dreifuss Muscular Dystrophy (AD-EDMD), Dilated Cardiomyopathy, Dunnigan type Familial Partial Lipodystrophy, Charcot Marie Tooth neuropathy, Hutchinson Gilford progeria syndrome, and others (Worman and Courvalin .2005; Young .2005). Emery Dreifuss muscular dystrophy (EDMD), Limb-girdle muscular dystrophy 1B (LGMD1B), dilated cardiomyopathy (DCM), and LMNA associated congenital muscular dystrophy(L-CMD) are the four forms of muscular dystrophies caused by about 74% of the mutations .Emery-Dreifuss muscular dystrophy (EDMD), the most severe and common Lamin A associated autosomal dominant muscular dystrophy, is characterized by progressive muscle loss and weakness, severe contractures of the elbows, posterior cervical, and ankles; conduction defects in the heart, and impairment in the distribution of humeroperoneal distribution (Bonne et al. 2000, Brodsky et al. 2000). In contrast, no contractures were detected in people with limb-girdle muscular dystrophy 1B (LGMD1B) (Muchir et al. 2000). The Lmna delK32/+ mouse model, which causes EDMD and L-CMD, has cardiac systolic dysfunction, ventricular dilatation, and dies young (Cattin et al. 2013). Lamin A/C regulates myonuclear mobility and localization in the periphery of myofibers, according to Roman et al., 2017. Due to the lack of laminA/C , muscle cells are more vulnerable to injury along with lower levels of lamin B1 . (Manilal et al.1999). When compared to wild type mice, myocardial histopathologies from male mice displayed aberrant elongated nuclei (Ostlund et al. 2001). LMNA^-/-^ mice also showed a lower fusion index in satellite cells due to lower amounts of myosin heavy chain genes (Myh1, Myh4, and Myl6h) than wild type mice (Cohen et al. 2013). This research group showed decrease in MHC immunostained cells reveal impairment in myotube development in LMNA^-/-^ cultures compared to WT. TGF/Smad signalling, MAPK signalling, Rb-MyoD signalling, and Wnt signalling all play key roles during muscle regeneration(Van Berlo et al. 2005, Muchir et al. 2007a).

Based on the previous studies , the pathogenesis of these muscular dystrophies can be explained by following mechanisms : i.nuclear fragility ,which might be generated by lower mechanical resistance (Broers, Ramaekers 2004; Paradisi, Djabali 2005; Vigouroux et al. 2001), ii. heterochromatin organisation (Vigouroux et al. 2001, McCord et al. 2013, Maraldi et al 2006; Scaffidi &Misteli 2005),iii. perturbation in cell proliferation or differentiation due to impairment in the interaction with regulatory factors (Capanni et al. 2005,Ivorra et al.2006).The myogenesis of lamin A linked muscular dystrophy mutants is governed by an unknown set of regulatory mechanisms. In cell culture, Holtzer et al. 1958, Kalderon and Gilula.1979 first demonstrated myoblast proliferation followed by migration and fusion. Mouse myoblast cell line C2C12 fills a significant niche in the gamut of cellular system accessible.Using this cell line we investigated biochemical changes behind the myoblast to myotube transition.In this study, we used two severe and well cited mutations of lamin A linked muscular dystrophy as mentioned in the database (www.umd.be/LMNA) : R453W and W514R ,to reveal molecular pathomechanisms that link different point muations of lamin A in muscle wasting phenotypes resulted by insufficient myoblast differentiation. To evaluate the differentiation potentiality of mutant LA cells, we first employed microscopy and immunoblotting approach. Gene regulatory factors responsible for myoblast to myotube commitment ,were studied using RNA sequencing. Systematic comparisons were done in order to have a complete understanding of a multifactorial differentiation process.

## Results

### Morphological assessment of mutant cells

Microscopical studies were performed on cells that had been allowed to differentiate for seven days. We found that mutant cells produced fewer myotubes per field (**Figure 1-a**). A closer look revealed the extent of the cell fusions and presented in **Table 1**. In mutant cells, we found that there was reduced multinucleation per myofiber with altered shape of the nucleus. We also observed that cells expressing mutant lamin A had smaller fibers. All of these findings were based on statistical analysis. These findings revealed that the mutant lamin A expressed cells had lower differential potentiality. Interestingly, LA R453W cells have lesser differentiation potential than LA W514 cells.

**Table 1:** Percentage of myotube formation per field.

| Sample Name<br>(Cell line : C2C12) | Day 7 (%) |
| --- | --- |
| WT LA | 72± 5 |
| R453W LA | 10± 1 |
| W514R LA | 15±1 |

**Figure 1:**
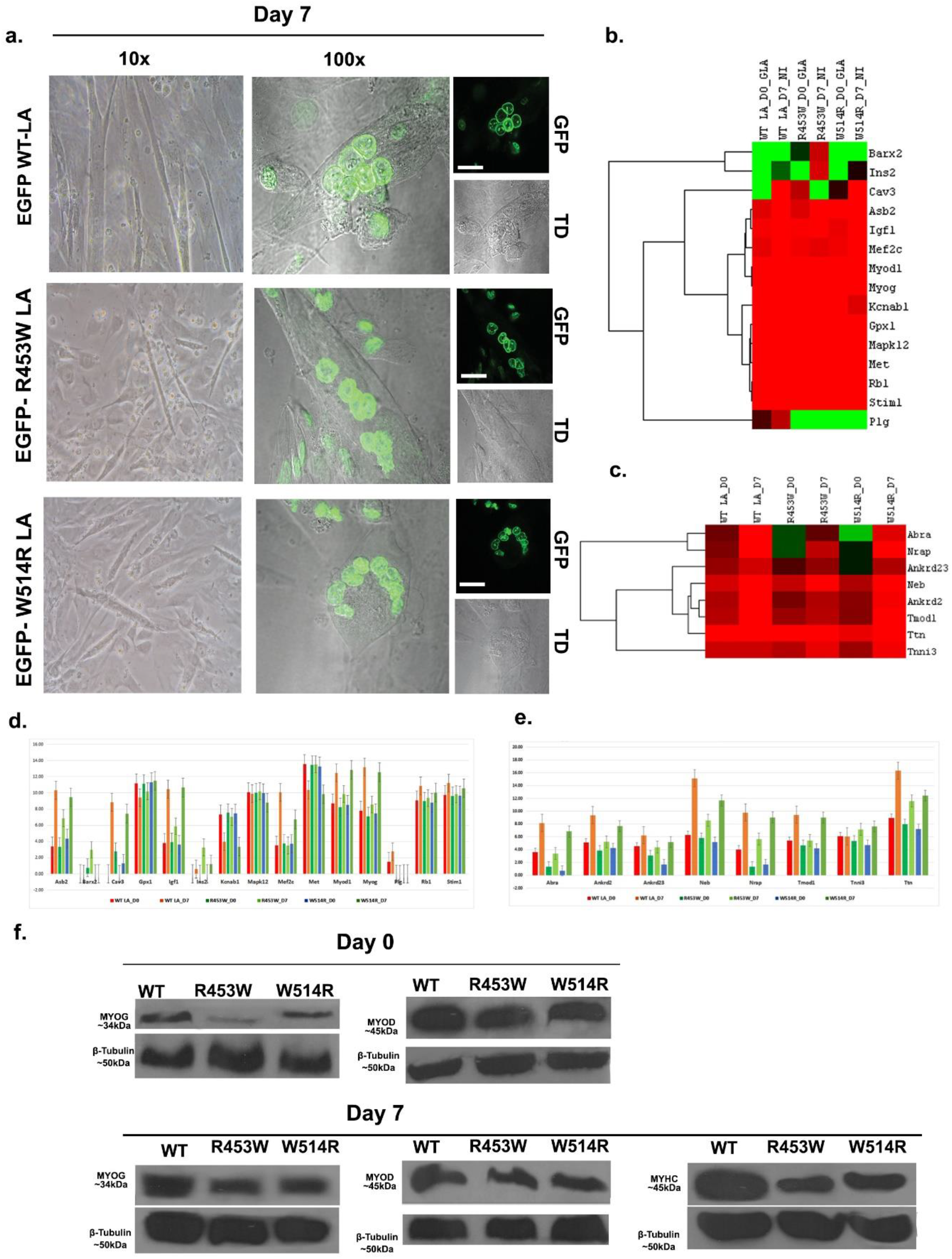
(a).After 7 days in culture, differentiation of myoblast into multinucleated cells is impaired due to muations. At an initial density of 1.6 × 10 ^5^ cells/well on 22 mm2 round cover glass in 35mm2 dish, cells were seeded and monitored under 10X and 100X objectives. Scale bar = 20 *µm*. (b), (d)present the heatmap and gene expression levels of Myoblast differentiation. (c),(e) are the heatmap and gene expression levels of Myofibril . (f). protein expression of Myo D, MYOG, and MyHC were monitored by western blotting. SDS PAGE was used to separate whole cell protein extracts after loading 35 µg proteins for immunoblotting with antibodies to β-tubulin, Myo D, MYOG, and MyHC. β-tubulin was used as loading control.

### Reduced levels of myogenic regulatory factors in mutant cells

In the myogenic regulation mechanism, there are number of mediators. MRFs (myogenic regulatory factors) have a coordinated effect during myogenesis (Weintraub et al. 1991, Rudnicki and Jaenisch 1995). When compared to the wild type, genomic analyses revealed that mutants have lower levels of expression of MRFs and other muscle-specific genes(**Figure 1-b,c,d,e**).

Immunoblotting was used to examine the protein level of differentiation markers . We observed significantly lower expression of Myo D,MyoG and MyHC in LA R453W, LA W514R cells than in WT LA cells (**Figure 1-f**). Overall, these findings revealed that mutations in lmna reduced differential potentiality of cells.

### Patterns of gene expression during the early stages of myogenic development

The gene expression reproducibility among the myoblast samples were demonstrated using principal component analysis, correlation matrix, and condition tree. 24995 genes were found with respect to three experimental conditions. From the hierarchical clustering analysis to reveal overlapping clusters of genes , we observed up-regulated and down-regulated genes were exclusively differently expressed in the undifferentiated myoblast cell population. To examine the distribution of upregulated and downregulated genes, the p value of Benjamini and Hochberg’s False Discovery Rate (FDR) <=0.05 and Fold Change Cutoff >=2.0 were used. When compared to the wild type LA, the volcano plot revealed that 318 genes were upregulated and 1342 genes were downregulated in LA R453W ,while 339 genes were upregulated and 958 genes were downregulated in LA W514R. The DEGs were then categorized into distinct functional groups (GO terms). A total of 105 GO terms and pathways were classified. The most statistically significant GO and pathways were classified into seven categories (**Figure 2**). According to the functions related to these genes , they are : Extracellular region (24 genes), signalling pathways (24 genes), epigenetic pathway (21 genes), gene expression regulation (14 genes), myofibril (8 genes), Ca+ ion transport (14 genes), and inflammatory response (13 genes) .

**Figure 2:**
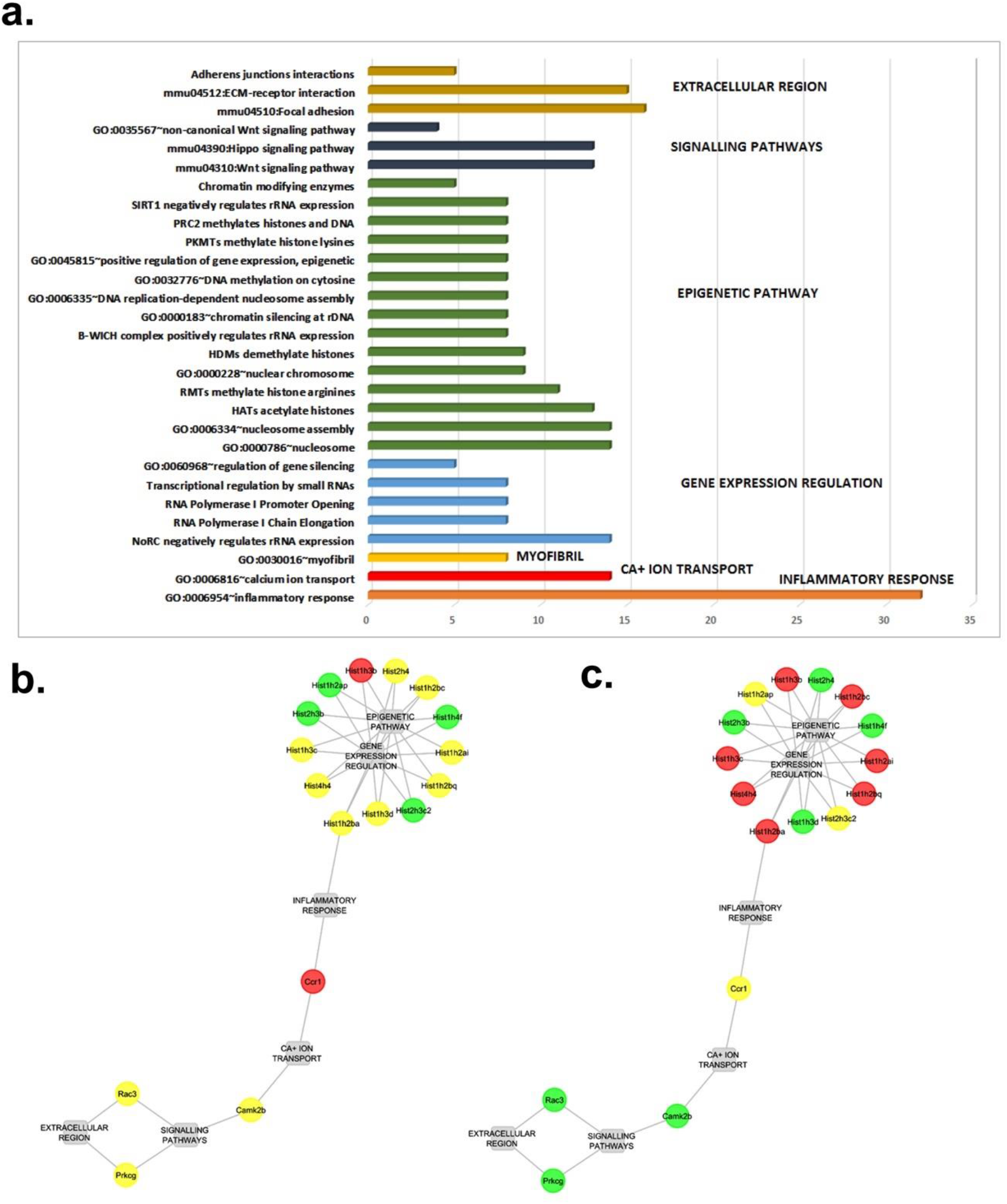
(a) Pathways that are dysregulated in the D0 state. (b), (c) Gene regulatory networks of R453W LA and W514R LA mutations .

### Patterns of gene expression during the myoblast differentiation process

Principle component analysis, correlation matrix, and condition tree were used to assess reproducibility among three samples of differentiation. The analysis of differentially expressed genes was done with a p value range of FDR<=0.05 and a fold change criterion of >= 2.0. In comparison to the wild type LA, we discovered that in LA R453W, 2145 genes were upregulated and 1933 genes were downregulated, while in LA W514R, 1789 genes were upregulated and 2075 genes were downregulated. Genes that were upregulated and downregulated were mapped to 707 GO terms and pathways, which were then assimilated into 14 functional groups (**Figure 3**). Extracellular region (186 genes), ribosome (98 genes), cytoskeleton (119 genes), signaling pathway (1042 genes), epigenetic pathway (114 genes), cell migration (180 genes), skeletal muscle myosin thick filament (13 genes), ion transport (14 genes), myoblast differentiation (173 genes), nuclear membrane (107 genes), cell division (30 genes), gene expression regulation(30 genes), myofibril (136 genes), Ca+ ion transport (42 genes) have been identified .

**Figure 3:**
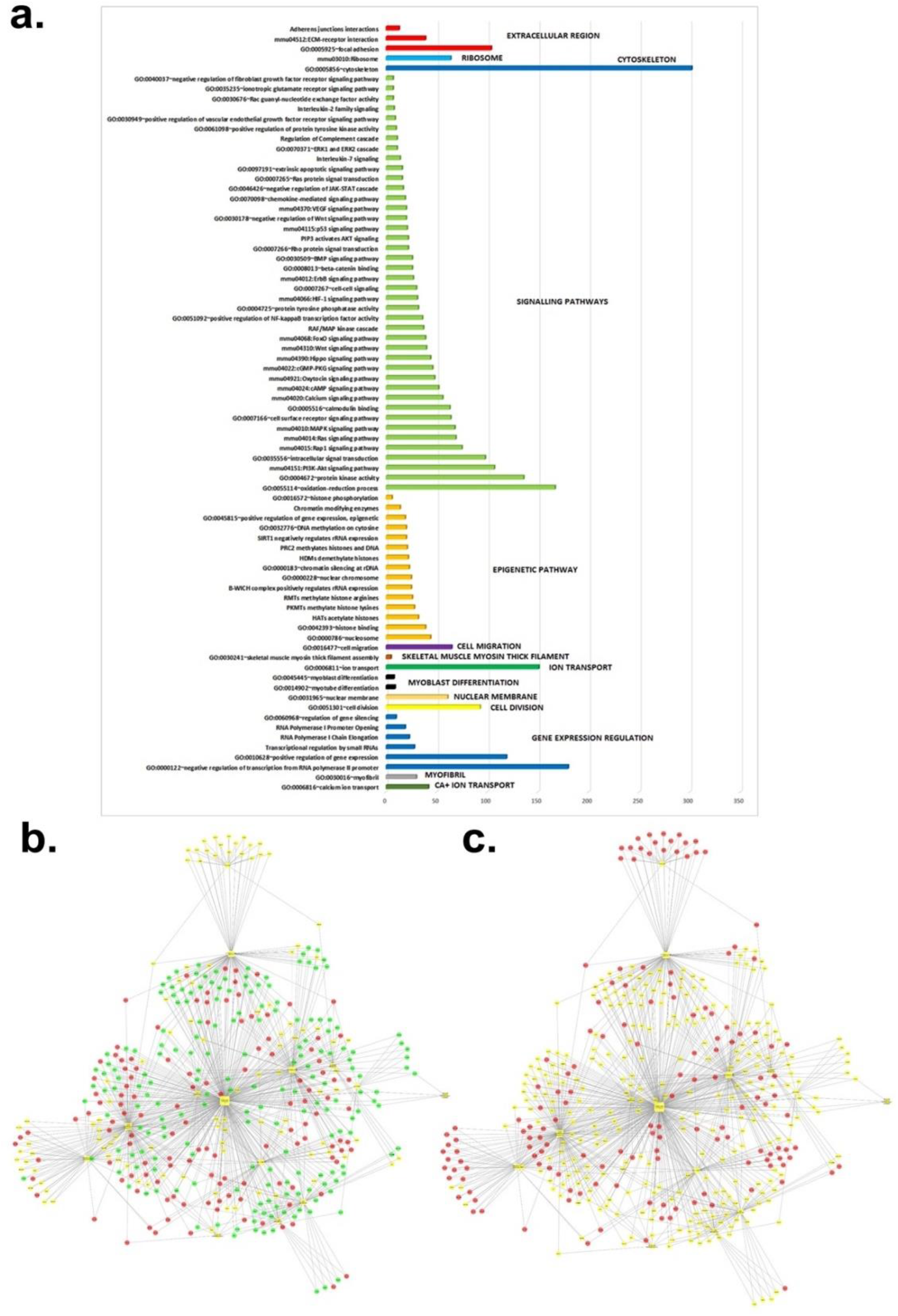
(a) Pathways that are dysregulated in the D7 state. (b) and (c) Gene regulatory networks of R453W LA and W514R LA mutations of day 7 state.

### Analysis of Signaling Pathway associations

Signaling cascades are generally responsible for satellite cell differentiation (Bentzinger et al. 2010). In particular, while comparing the differentiation programme of mutants to the differentiation programme of wild type LA, we discovered 40 dysregulated (**Figure 3**) signaling pathways.

Interestingly, we found Wnt and Hippo signaling pathways are common dysregulated significant signaling pathways in each of the 6 experimental conditions and their role as a modifier of muscle differentiation already validated previously (Maltzahn et al.2012). We observed depression of Wnt 6 gene in the differentiated state of mutant cells, which is a primary regulator of Wnt signaling cascade/Hippo signaling cascade. Our study also uncovered reduced expression of Nfatc2, Tcf7L2, Lef1,Gli2 transcription factors (**Figure 4**). Furthermore, at the differentiation stage, mutant cells had a high amount of Apc2 gene expression. The Apc2 gene contains a binding site for β-catenin and has the ability to degrade it, making it an important component of the Wnt pathway. As a result, a high amount of Apc2 indicates that β-catenin transcriptional activity may be inhibited (Kennell et al.2009).Resultantly, the Wnt signaling pathway/Hippo signaling pathway may have been downregulated in mutant differentiated cells.

**Figure 4:**
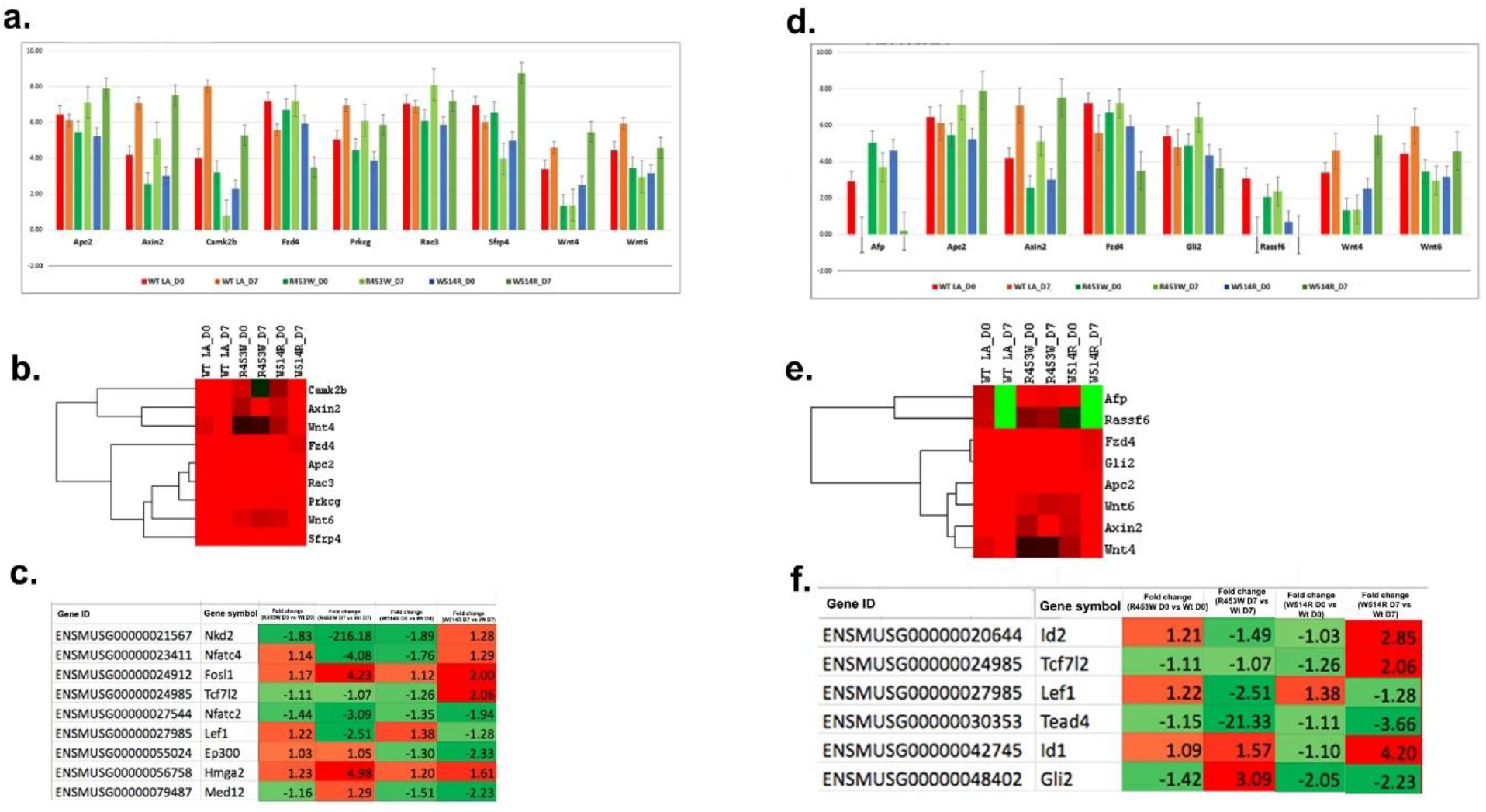
(a),(b).Alteration of expression of gene products of Wnt pathway in two mutant samples were expressed as gene expression profile and heatmap including undifferentiated and differentiated stages.(c).The transcription factor level of the Wnt pathway in all samples in the D0 and D7 conditions. (d),(e). The gene expression profile and heatmap of the Hippo pathway in three samples at undifferentiated and differentiated stages (f) represents the transcription factor levels of Hippo pathway in all samples.

### Changes in Ca+ ion transport

Calcium ion transport regulates the expression of MRFs during myogenic differentiation through influencing Ca ion dependent kinases and phosphatases (Tomczak et al.2004). From GO analysis of Ca ion transport pathway , we observed 14 genes were dysregulated in the undifferentiated stage and 42 genes were dysregulated in the differentiated stage. When comparing LA R453W cells to LA W514R cells in both undifferentiated and differentiated stages, we detected higher downregulation of Ca ion transport related genes such as Atp2B4,Atp2a1, Camk2b,Cacna1c, Nalcn, and Ryr3 compared to wild type LA cells (**Figure 5**).

**Figure 5:**
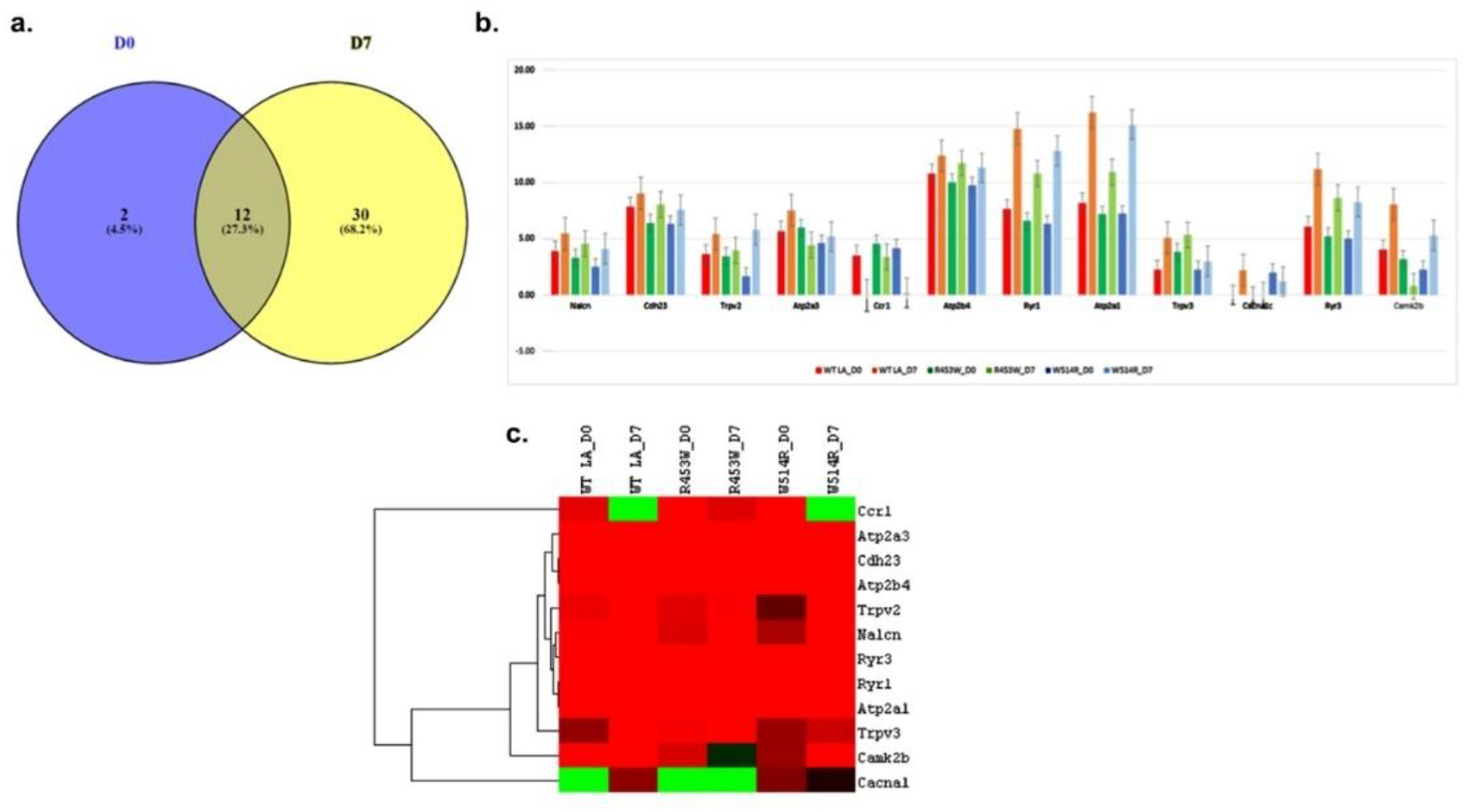
(a).Between D0 and D7 conditions Venn diagrams illustrate shared genes in the Ca+ ion transport system. (b),(c) Gene expression levels and a heatmap of the Ca ion transport pathway.

### Epigenetic regulation in differentiation

In muscle differentiation, epigenetic networks give an instructional signal (Perdiguero et al. 2009). Previous research has shown that heterochromatic detachments from the nuclear lamina occur in mutants of lamin A/C during differentiation (Perovanovic et al.2016,2018). When comparing mutants in the differentiation stage to mutants in the undifferentiation state, we found that mutants in the differentiation state had higher upregulation and downregulation of genes. We discovered 20 genes dysregulated in the undifferentiated stage and 114 genes dysregulated in the differentiated stage using GO analysis of epigenetics(**Figure 6-a,b**). Surprisingly, from the common mis -expressed genes analysis in genomic profiling revealed that, the Kdm7a gene has a low expression level. Eraser protein Kdm7a is a demethylase to demethylate the H3K27me3 restrictive chromatin mark. On the other hand, we discovered a low level of the Ogt gene, which is an O-GlcNAc transferase. According to Yang et al.2002, the chromatin repressor mSin3A interacts with Ogt and prevents O-GlcNac modification in the activation of RNA pol II enzyme for active transcription. Furthermore, the immunoblot study revealed a high protein expression level of repressive chromatin mark (H3K27me3) in both LA R453W and LA W514R cells at differentiated stages(**Figure 6-d**).

**Figure 6:**
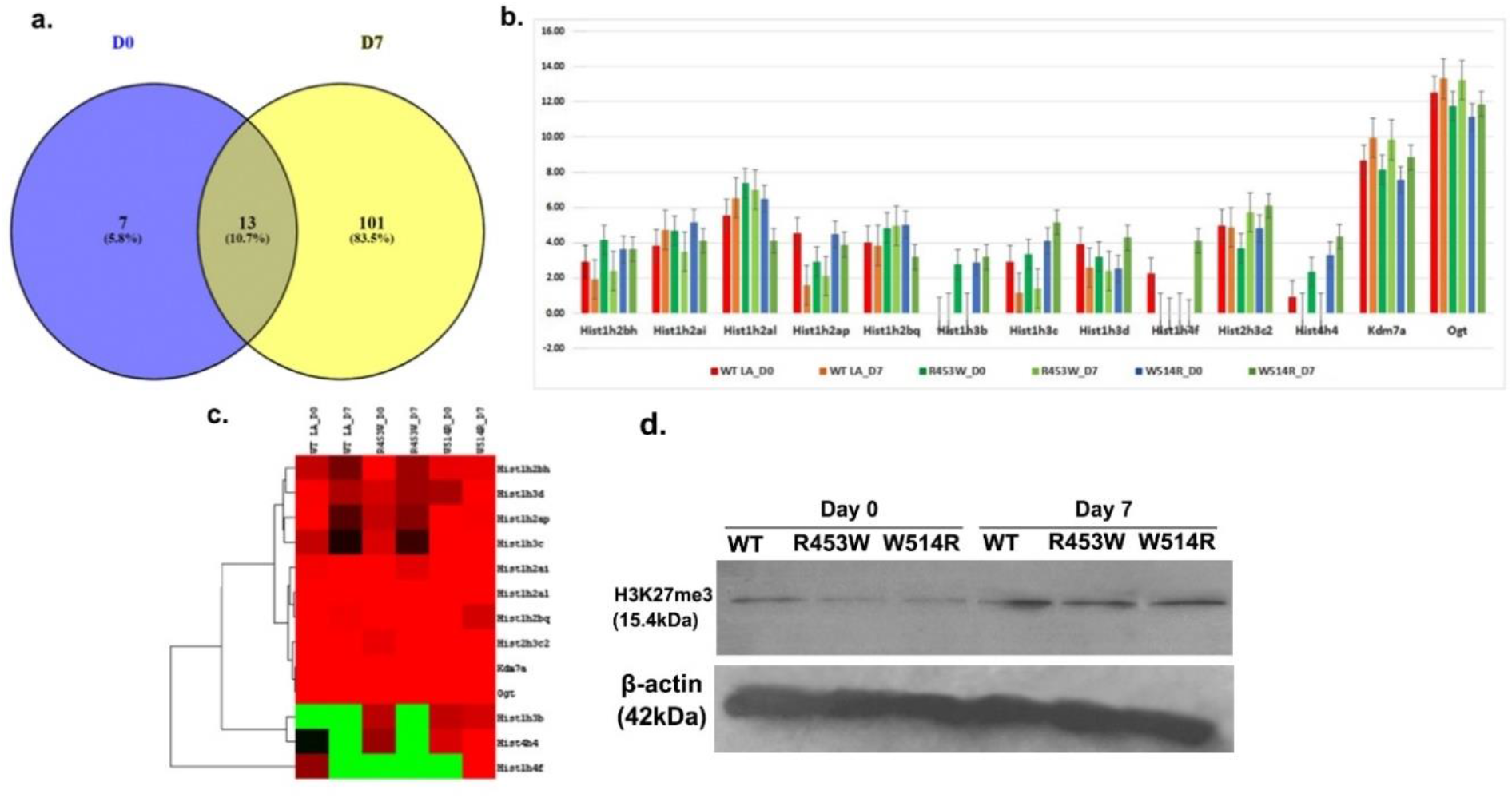
(a).Genes in the epigenetic pathway that are common in D0 and D7 conditions. (b),(c) represent gene expression levels and epigenetic pathway heatmap. (d)H3K27me3 protein expression levels after immunoblotting with antibodies to β-actin (control) and H3K27me3.

### Difference in the expression of focal adhesion proteins and cell motility in mutants

Focal adhesion proteins make permissive environment for the myoblasts fusion by influencing the cell migration. Members of this protein family act as avenues for communicating with extracellular matrix components on the outside of the cell as well as with cytoskeletal components in the cell interior. We assessed changes in paxillin protein distribution in lamin A Wt and mutant cells at day 0 phase. This analysis revealed less recruitment of paxillin at the periphery of mutant cells. At the genome wide level we found unbalanced levels of GO pathway focal adhesion with the downregulation of genes, like – Cav3,Cd36, Cdh11,Col24a1,Col6a2, itga11,Mylk3,Prkcg,Vwf.So, it is tempting to speculate impairment of the cell migration in mutants, which may contribute to the disease phenotype. In line with this hypothesis ,our observation from wound healing assay showed less number of cells at the wound closure area in mutant cell model system. This part will be elaborated later. Results of this part are shown in **Figure 8**.

## Discussion

Our findings postulate a physiologically important role of lamin as a modifier of muscle differentiation fate in lamin A associated muscular dystrophies and eventually the impaired muscle function. To understand the muscle differentiation programme in muscular dystrophy mutations, our study is a cumulative effect of parallel and stage specific events of impaired heterochromatinization, reprogramming of epigenetic mechanisms, destabilized transcriptome profiling, poor induction of signaling molecules, and myogenesis effectors.

Our results revealed that LA R453W cells exhibited lower differentiating potential than LA W514 R cells, indicating that the R453W mutation has a devastating effect on muscle function recapitulation. Our findings shed light on the molecular regulators involved in the differentiation process in lamin A-related skeletal muscle degeneration. We found a constant drop in MyoD level in mutant LA cells in a time course dependent manner, which hindered the fusing of myoblast cells in the creation of myotube, based on immunoblot and RNA sequencing data. MyoD is a vital myogenic regulatory factor and it’s expression starts before the differentiation and has a critical function also during the entire differentiation stages (Yin et al.2013). In both undifferentiated and differentiated stages, mutant cells have lower amounts of myogenin. Myogenin is also a critical component in myogenesis, as its presence is sufficient to induce myoblast fusion (Hasty et al. 1993, Rawls et al. 1995,Venuti et al. 1995). In mutant LA cells, however, we found a lower amount of the terminal differentiation marker MyHC when compared to wild type LA cells. As MyHC is a component of myofibril proteins (Klont et al.1998, Ryu et al.2005).Diminished expression of MyHC by immunoblot analysis assign an alignment with the decreased expression of myofibril genes (Abra, Ankrd2, Ankrd23, neb, Tmod1, Ttn) in mutant LA cells in the differentiation stage (Figure.1-c,e). In addition, we found lower expression of Ca+ ion transport related genes, which are involved in the disorganization of myofibrils, as described by Zvaritch et al.2007. Myofibrils are made up of contractile muscle units called myofilaments, which are complexes of actin and myosin interaction. MyHC has a head, neck, and rod domain. The head domain binds to actin, while the neck domain binds to myosin. As a result of our aforesaid findings, it appears that a defective differentiation programme may result in muscle wasting, compromising muscle contractile function in lamin A linked muscular dystrophy due to impediment in myofibril formation.

With the association of histones, Lamin A tethers heterochromatin at the nuclear periphery, which tends to detach from lamina upon myoblast to myotube transition (Mattout et al. 2007, Taniura et al.1995, Perovanovic et al. 2016,2018). Importantly, as the integrity of the lamina was disrupted, we found more upregulated and downregulated genes in the differentiation of mutant LA cells (**Supplementary Figure.1**). (e). Furthermore, we discovered a significant increase in histone repressive marks, such as H3K27me3, in the differentiation stage of mutants using our immunoblot method, which was examined for the first time. In support of this, gene expression profiling revealed that mutants have lower levels of Kdm7a and Kdm4c, which are demethylases of H3K27me3 and H3K9me3 modifications, respectively. The plausible explanation is based on the interaction of lamin A/C with muscle gene specific promoters in myoblasts, which decreases as differentiation progresses, as discovered by Athar et al.2015. In normal cells, they also discovered an increase in the population of active histone marks on the MyoG and MyoD1 gene loci. Lu et al. (2000) and Fu et al. (2013) discovered that muscle-specific genes are expressed during differentiation due to histone acetyltransferase enrichment. To support this,our bioinformatic analysis showed downregulation of Rb1gene in the mutants. Rb1 helps in MyoD mediated transcription of myogenic target genes by relinquishing HDAC1 from MyoD(Magenta et al. 2003, Mal et al. 2003, 2001).Thus, it may be a contributing factor to the acetylation of MyoD . Interestingly, in human muscle biopsy data, upregulation of four genes (CREBBP,CRI-1,NAP1L1 and EP300) were observed (Bakay et al. 2006). In support of this, we identified downregulation of Rb1, Ep300,Creb 5 ,that indicates attenuation in the acetylation of MyoD in the differentiated states of mutant cells. Decreased level of Rb1 gene also represents less recruitment of it to the lamina with the disruption of Rb1/HDAC complex formation and repression of Rb1/MyoD pathway as well as Rb1/E2F pathway. Overall, it validates chromatin reconfiguration in mutants due to changes in lamina structure.

We discovered that the Wnt and Hippo pathways are frequently dysregulated pathways in mutant cells at both the undifferentiated and differentiated stages, based on RNA sequencing profiling. Wnt and Hippo signalling cascades were shown to be downregulated in mutant LA cells in both the undifferentiated and differentiated stages, indicating that these signaling pathways are important in the differentiation process (Singh et al. 2012). Wnt6 gene downregulation and Apc2 gene overexpression were found in our RNA sequencing results. Lavery et al.2008 and Geetha-Loganathan et al. 2005 found that Wnt6 modulates the canonical Wnt/catenin signaling pathway and positively affects the production of Myogenin, Myf5, and MyHC. Downregulation of the Wnt signaling pathway, as well as the concentration of repressive chromatin marks in the differentiated state of mutants, could have influenced downstream effectors such as, transcription factors and cofactors. In line with this, we detected significant downregulation of the following transcription factors (mentioned in Figure 7), which directly or indirectly affect muscle differentiation in mutants, as indicated by a plethora of evidences : MEF2A (Kaushal et al. 1994), MEF2C (Lin et al. 1997), Rb1 (Novitch et al. 1996, Schneider et al.1994), Tead 4 (Benhaddou et al. 2012), Asb5 (Ehrlich et al. 2020), Asb 15 (Tee et al. 2010), Bcl6 (Kumagai et al. 1999), NR4A1 (Pan et al.2019), Gli (Lin et al. 2014), Nfatc2 (Armand et al.2008), Clock (Andrewset al. 2010).According to earlier studies, downregulated transcription factors are critical components of numerous signaling pathways, several of which were shown to be dysregulated in our investigation. Future study into how these pathways interact through co-ordinated molecular interactions will provide a clearer picture of the defective differentiation programme in lamin A-associated muscular dystrophy. Moreover, multiple layers of redundancy among the pathways could slow the progression of the disease in form of relatively slow muscle loss phenotype . This also predicts later onset of dystrophic phenotype due to Lmna related muscular dystrophies.

**Figure 7:**
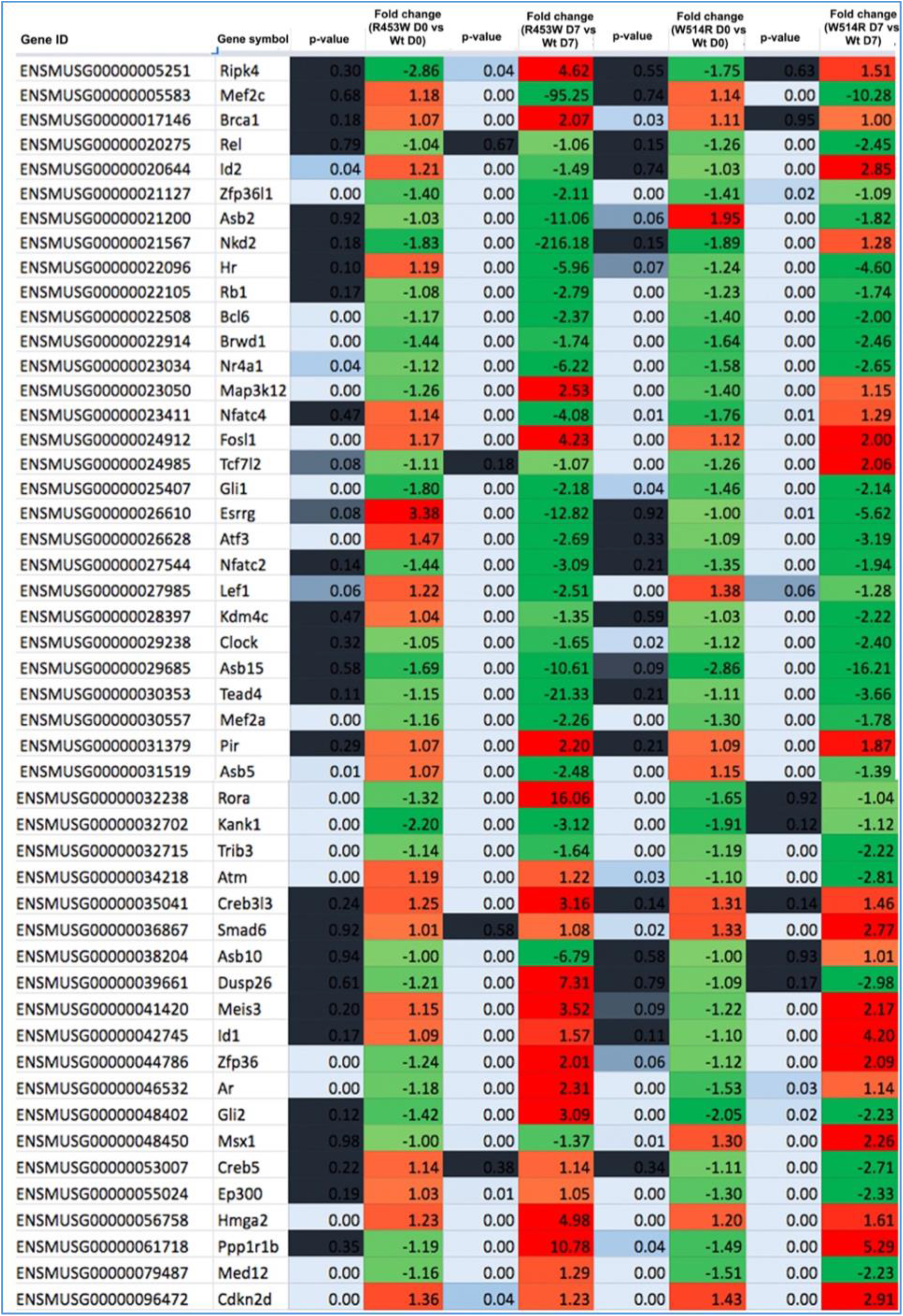
List of total transcription factors of R453W LA and W514R LA after substituting expression levels of WT LA in D 0 and D7 states . Green color indicates down regulation and red color indicates up regulation.

**Figure 8:**
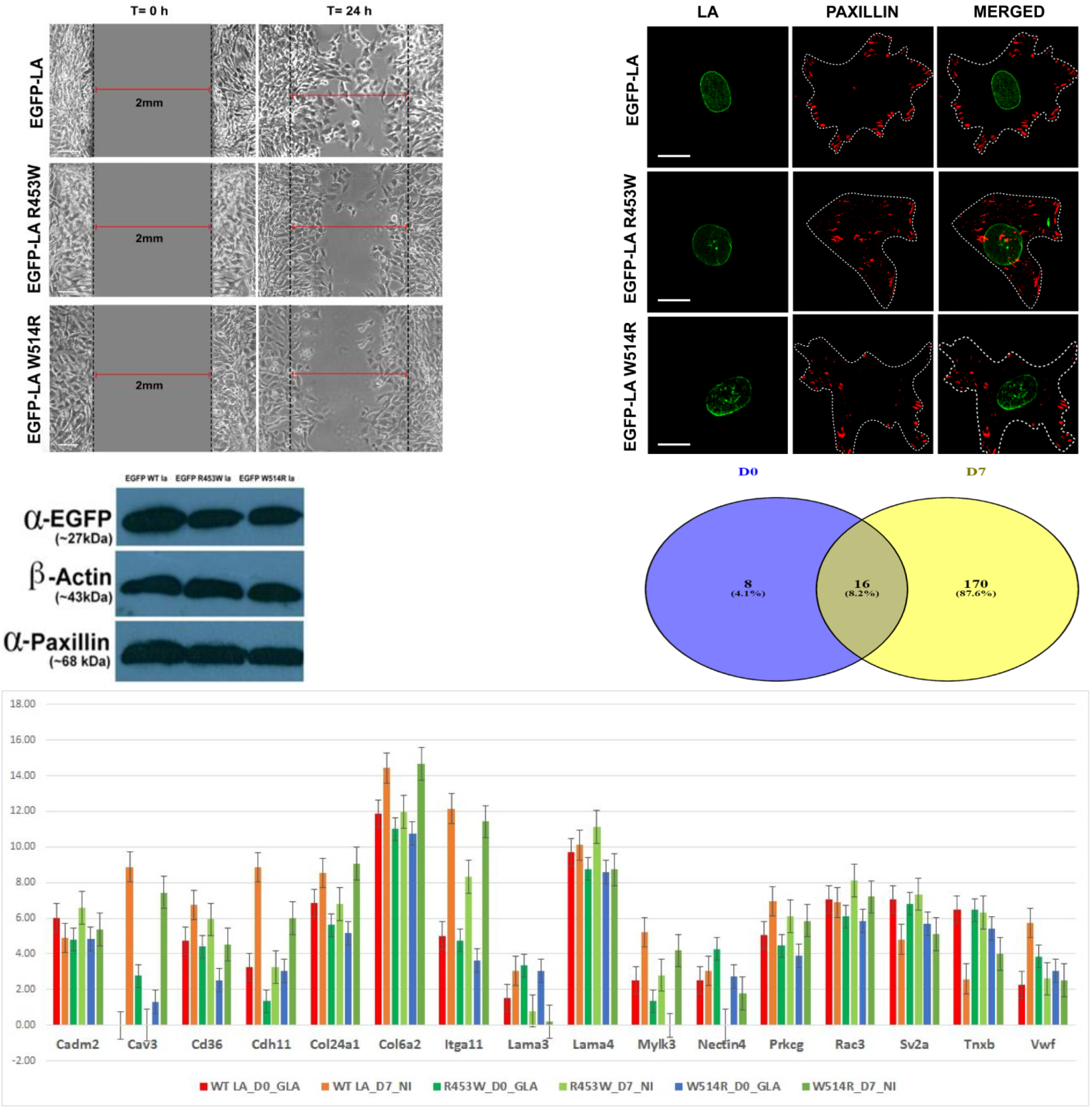
**(This figure is in the process of being updated)** **Mutaions in lamin A lead to perturbation in migration** , **consistent with downregulation of components of focal adhesion complexes**. (a). The wound healing assay reveals differences in cell migration in mutant cells.(b). R453W LA, W514R LA cells stained for paxillin and confocal images illustrates less deposition of paxillin in the cell periphery.(c). Changes in GFP lamin A, paxillin,β-actin were confirmed byimmunoblot for wild type and mutant day 0 phase cells.(d).Genes in the “extracellular region” that are common in D0 and D7 conditions.(e). Gene expression levels of “extracellular region”.

## Methods

### Stable cell line preparation

After Lipofectamine 2000 transfection of mouse myoblast cell line C2C12 with EGFP-LA, EGFP-R453W, and EGFP-W514R constructs (constructs were generated in Bera et al.2004, Dutta et al.2018,2022) , selection antibiotic G418 was standardized to 850 µg to select positive GFP clones. For 3-4 weeks, maintenance medium that supplemented with G418 selection antibiotic ,was replaced on a daily basis.

### Cell culture

The stable transfected mouse myoblast cell line C2C12 (ATCC, Manassas, VA, USA) was cultured in DMEM (Life Technologies) high glucose media with 10% fetal bovine serum, 100 units/ml Penicillin, and 100 g/ml Streptomycin at 37°C in a 5% CO2 incubator. The medium was changed to DMEM medium with 2 % horse serum, 100 units/ml penicillin, and 100g/ml streptomycin for differentiation initiation when the cell confluency reached 80 percent. Fresh differentiation media was added every two days and the medium was allowed to differentiate for seven days.

### Cell lysate preparation &protein quantification

Scraper was used to lift cells for cell lysate production. 7M Urea buffer supplemented with Mammalian Protein Extraction Reagent (M-PER) (Thermo Scientific, USA) and 1X protease inhibitor cocktail was used to lyse the cells, which were then sonicated using a UP 200s (ultraschallprozessor). After that, it was acetone precipitated overnight at -20 C, then centrifuged for 15 minutes at 13000 rpm. To get rid of the acetone, the pellets were air dried. To disintegrate the pellet, 50 µl of 7M urea buffer of above composition was added. The Bradford reagent was used to determine the protein concentration, with BSA as the standard.

### Immunostaining and microscopy

Cells were fixed in 4 % paraformaldehyde for 10 minutes before and after differentiation (D7), permeabilized with 0.5 percent (v/v) Triton X-100 in PBS for 5 minutes, then blocked with 5% (v/v) normal goat serum in PBS. VECTASHIELD mounting media (Vector Laboratories, Peterborough, UK) was used to mount coverslips on cover slides. Images of multinucleated cells (n=150) were captured using an NIKON Inverted Research Eclipse TiE Laser Scanning Confocal/NSIM microscope with a Galvano mode NIKON A1 RMP detector and Plan Apochromat VC 100x oil DIC N2 / 1.40 135 NA /1.515 RI objective and an extra 4x digital zoom. For the green channel, a multi-line Argon-Krypton laser (ex-457/488/561 nm) operating at 3% was used. A 0.25 m step size was retained for Z stacks. By using the 10X objective of a Nikon TS100 Inverted Phase Contrast Microscope, 200 cells (containing mono and multinucleated cells) were examined to studying the differentiation potentiality of each sample (WT LA, Mutant LAs).

### Western blot

On a 7 % polyacrylamide gel, 35 µg protein samples from days 0 and 7 of each sample were loaded. To transfer separated proteins onto a PVDF membrane, semi-dry transfer method was used. After that, the PVDF membrane was blocked with PBS solution containing 5% nonfat dry milk for 1 hour before being incubated with primary antibodies : MyoD (ab203383) at 1:100,Myogenin (ab1835) at 1:250,MyHC (MAB4470) at 1:500 and actin (Sigma–Aldrich, A5316) at 1:800. A secondary antibody linked to horseradish peroxidase (HRP) was used at a dilution of 1: 1000. Following that, the membrane was developed using (Thermo Fisher Scientific Inc) Super Signal West Pico reagent. Post that, densitometry analysis was performed to measure signals using Image J (Version 1.48) as described by Schneider et al. 2012.

### Sequencing and analysis of RNA

Total RNA was isolated using a Qiagen RNA isolation kit. A spin column-based RNA isolation kit was used to isolate total RNA (Qiagen). 50 µg of total RNA was used as the starting material for whole transcriptome library preparation after RNA was quantified using a NanoDrop UV-VIS spectrophotometer at 260nm. After running the Bio analyser, the cDNA library was sequenced on the Ion PGM™ system. This system was first initialized with dNTP reagents of Ion Hi-Q sequencing 200 kit.

### Analysis of bioinformatics

Raw data of the samples were obtained in FASTQ format from the H+ Torrent server, with a total read of 7.5 million and a mean read length of 100 bp. FASTQ files were processed with Ion Torrent’s built-in WT pipeline, and reads were aligned to the mm10 reference genome using the TopHat aligner as an annotation model. Cluster 3.0 programme was used to plot Condition tree, PCA (Principle Component Analysis), and Correlation Matrix to find out if there was any link between the samples. The greater the similarity in gene expression between samples, the closer they are. The purpose of a correlation matrix is to figure out how many genes were expressed in both samples. A higher correlation score indicates that more genes are expressed in both samples. The software PartekFlow™ (Partek Inc, MO, USA) was used to find differentially expressed genes (DEG s) in wild type and mutant Lamin A s. Statistically significant differentially expressed transcripts with FDR score of <= 0.05, p value of < 0.01 and fold change of -2 to +2 were subjected to GO and Pathway enrichment with the DAVID tool (Ashburner et al. 2006, Huang et al. 2008). The Pathreg algorithm was used to examine the gene regulatory network derived from GO terms, as well as the differentially expressed genes contained them. Shannon et al. (2003) used Cytoscape v2.8.2 to find critical nodes and edges that could be reflective of gene regulation alterations caused by mutations.

### Wound healing assay

Approximately 2х10^5^ cells per 6 well dish were seeded the night prior to wound formation. Confluent monolayers were then scratched with a 2µl pipette tip . Cell debris was removed via PBS washing and replace with fresh medium. Consistent pressure and pipette tip angle should be kept to make the wound consistent. Images were taken in bright field using 10X objective lens after 24 hour.Analysis of wound closure computed using Image J.

## Acknowledgements

The authors thank Ashish K.George (Thermo Scientific),Mr. Saikat Mukherjee (Crystallography and Molecular Biology Divison, Saha Institute of Nuclear Physics) for the technical support in RNA sequencing experiment. We sincerely acknowledge contributions of SERB-DST and BARD project,DAE for partly funded the research.

## Author Contributions

**SD**, conceptualization,resources,methodology,investigation, data curation,formal analysis and interpretation,writing the manuscript.**MV**, supervision,resources, analysis and writing the manuscript.All authors have approved the final draft of the manuscript.

## Conflict of interst

The authors declare no conflict of interest.

---

**Figure supplementary 1:**
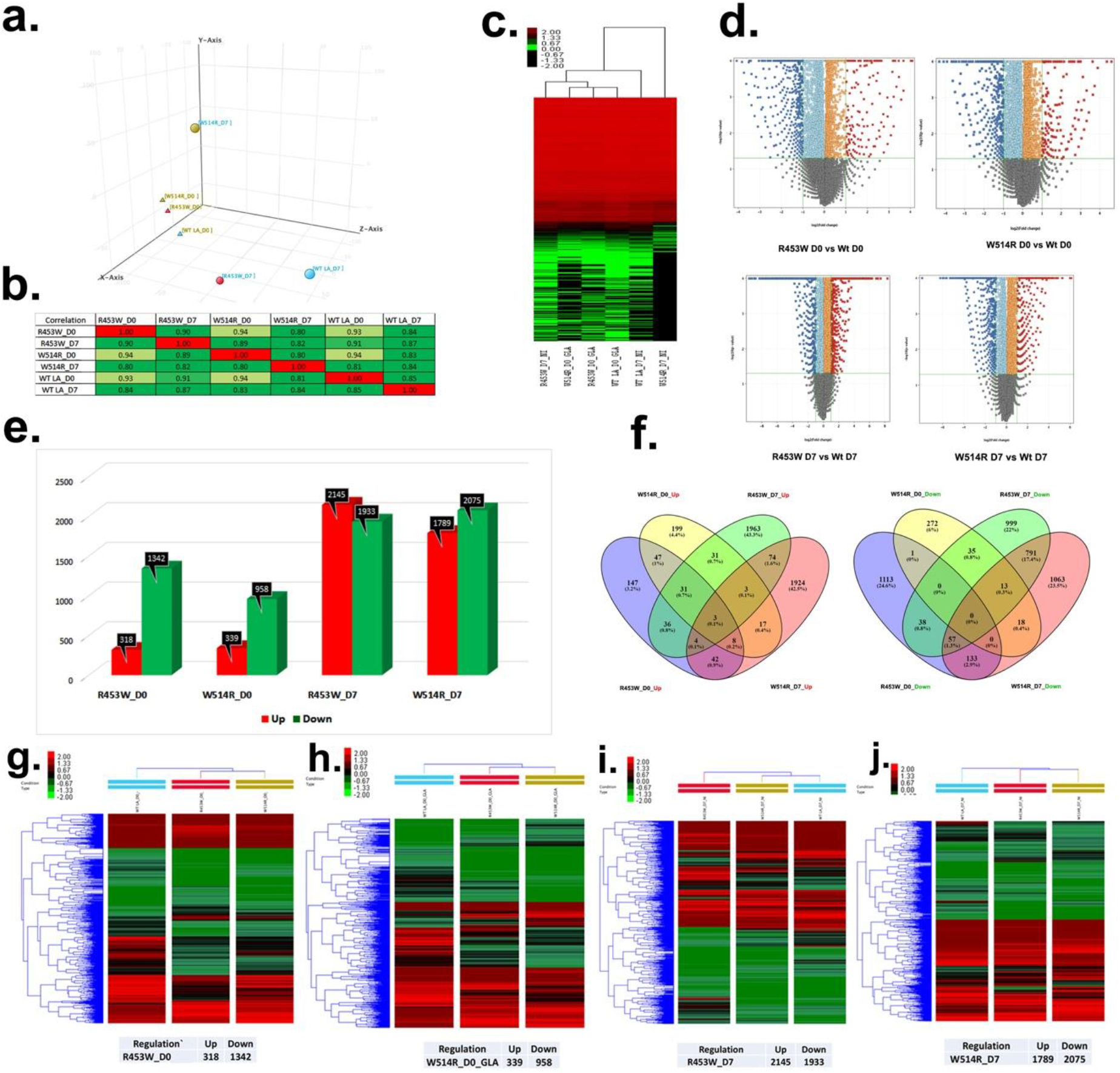
(a), (b), and (c) are qualitative assessments of the Deep Sequenced Transcriptome: PCA analysis, correlation plot, and condition tree, respectively, from day 0 and day 7 of three samples.(d) Volcano plots of each different mutant sample following wild type sample substitution. (e) The histogram depicts the up and down-regulated genes in each sample. (f) Venn diagrams of R453W LA day 0 against W514R LA day 0 and R453W LA day 7 vs W514R LA day 7 . (g), (h), (i) and (j) Cuff Diff pipeline was used for unsupervised hierarchical clustering to show differentially expressed transcripts. Fold change criterion of >=2.0 was set, as well as a student t-test p value of 0.05. Red denotes upregulation, black shows no regulation, and green indicates downregulation in the clusters.

## Notes

### Competing Interest Statement

The authors have declared no competing interest.

